# Development of resident and migratory three-spined stickleback, *Gasterosteus aculeatus*

**DOI:** 10.1101/2023.12.05.570243

**Authors:** Megan Barnes, Lisa Chakrabarti, Andrew MacColl

## Abstract

The three-spined stickleback (*Gasterosteus aculeatus)* is a teleost fish and a model organism in evolutionary ecology, useful for both laboratory and natural experiments. It is especially valued for the substantial intraspecific variation in morphology, behaviour and genetics. Classic work of Swarup (1958) [1] has described the development in the laboratory of embryos from a single freshwater population, but this was carried out at higher temperature than many stickleback would encounter in the wild and variation between populations was not addressed. Here we describe the development of embryos from two sympatric, saltwater ecotypes of stickleback from North Uist, Scotland raised at 14°C, the approximate temperature of North Uist lochs in the breeding season. The two ecotypes were (a) a large, migratory form in which the adults are completely plated with bony armour and (b) a smaller, low-plated form that is resident year-round in saltwater lagoons. By monitoring embryos every 24-hours post fertilisation, important characteristics of development were observed and photographed to provide a reference for North Uist ecotypes at this temperature. Hatching success was greater than 85% and did not differ between resident and migratory stickleback, but migratory eggs hatched significantly earlier than the resident ecotype. Our work provides a framework that can now be used to compare stickleback populations that may also grow in distinct environmental conditions, to help understand the breadth of normal developmental features and to characterise abnormal development.

## Introduction

The three-spined stickleback has been increasingly studied and raised in aquaria [2]. The repeated adaptation of oceanic stickleback to freshwater make it an attractive model to investigate parallel evolution [3]. As such, the three-spined stickleback has become an important model in evolutionary genomics, with both laboratory and natural experiments widely reported. Despite much recent research on variation in the morphology, behaviour and genomics of stickleback, there is a paucity of work describing variation in the development of stickleback from fertilisation to hatching. Swarup 1958 described in detail the key stages in development of the three-spined stickleback, building upon Vrat, 1949 [4] and Kuntz & Radcliffe, 1917 [5]. However, Swarup’s embryos developed in the lab at 18-19°C, a relatively high temperature for stickleback, and did not consider the possibility of variation between populations. Given the wide distribution of stickleback across the Northern Hemisphere encompassing large variation in environmental variables and therefore stickleback morphology, it is useful to quantify development of the stickleback at lower temperatures, and to compare contrasting ecotypes.

On North Uist (Western Isles, Scotland), there are two morphologically distinct ecotypes that occur sympatrically in saltwater: migratory and lagoon resident [6]. Migratory stickleback are large and completely plated, spending most of their lives at sea but migrating to brackish water to spawn, whereas smaller, low plated lagoon resident fish live permanently in coastal lagoons. Resident stickleback lay smaller clutches than migratory, and these consist of larger eggs. High levels of reproductive isolation are maintained between the ecotypes [6]. Here we compare the development of migratory and resident stickleback from North Uist at a temperature naturally experienced by these populations, and provide reference photographs for future work.

## Methods

To compare development in sympatric stickleback ecotypes at a physiologically relevant temperature (14°C) and provide a photographic resource for future studies, we monitored the development of lagoon resident and migratory stickleback from North Uist, starting at fertilisation through to hatching. Key features were noted every 24-hours and coloured photographs taken to record the key stages. Hatching success and time to hatching were also recorded for both ecotypes.

Fieldwork was conducted on North Uist between 30^th^ April and 19^th^ May 2023. To collect breeding wild stickleback of both sexes and ecotypes, mesh traps were left in Loch an Duin (57.64245, - 7.209207), a saltwater lagoon in the North-East of the island, for 24-hours. Migratory and lagoon resident crosses were made, following standard procedures [7], by squeezing eggs from gravid, euthanised females into small petri dishes and mixing them with testes from euthanised reproductive males. Fish were euthanised with an overdose of tricaine methanesulfonate (400mg/L) followed by destruction of the brain in accordance with Schedule One of UK Home Office regulations. After fertilisation, eggs were just covered in sterile water. Egg number per clutch was kept between 20 and 30 where possible. Six families per ecotype were made using different males and females for each, and each was raised in a separate petri dish. Embryos were maintained at 14°C, the approximate water temperature of North Uist Lochs in April, in an incubator (ICT-P Falc, portable mini incubator) where air could fully circulate. Every ∼24-hours post-fertilisation, petri dishes were briefly taken out of the incubator and media carefully removed with a Pasteur pipette and then replaced with fresh, temperature-matched media. Each dish was observed under a dissecting microscope (Olympus SZ61) and key features of developing embryos noted, as well as photographs taken. Embryo development was also scored based on Swarup 1958. This was repeated daily, until two days after the first hatched stickleback fry was observed.

Hatching success was calculated for each clutch of migratory and lagoon resident ecotypes separately as percentage of successfully hatched fry, and the number of days until the first appearance of a fully hatched fry was noted. Hatching success was compared between ecotypes using a binomial GLM with logit link. In one clutch, 17 unfertilised eggs were observed 48-hours post-fertilisation; this was likely due to hardening of the eggs in the female reproductive tract which can be common in stickleback [8]. In this case hatching success was recalculated as the number of successfully hatched fry per total number of fertilised eggs, and this value was used in the model.

Time to hatch (days) was compared between migratory and lagoon resident stickleback using a t-test. To assess differences in development between migratory and lagoon resident stickleback, the score based on Swarup’s 1958 taxonomy was plotted at each 24-hour timepoint for both ecotypes. We tested for differences between ecotypes in the development score over multiple days using two-sample Wilcoxon rank sum tests, and used the false discovery rate (FDR) method to control for type 1 errors [9]. An FDR of 0.10 was used. All analysis was done in R Studio.

## Results

The number of eggs selected per clutch averaged 23.6, with means of 27 and 20.2 eggs for migratory and resident clutches respectively (in total, clutches are larger than this, especially for migratory fish). There was no difference in hatching success between ecotypes (Wald X^2^_1_= 0.67, P = 0.41; Table 1, Figure 1a), with hatching success averaging 85.0% in migratory stickleback and 92.7% in lagoon resident fish (Figure 1a). The number of days until appearance of the first hatched fry was significantly earlier in migratory than lagoon resident stickleback (t.test: t = -5.58, df = 9.49, p = 0.00028; Figure 1b), with migratory fish hatching after 11.7 days on average (clutches hatched on days 11 or 12) and resident fish after 13.2 days (clutches hatched on day 13 or 14).

**Table 1:** Wilcoxon Rank Sum test results for comparisons between migratory and resident stickleback development scores at each day ranked in order of increasing p value, with multiple comparisons testing of results. FDR set to 0.10.

| Day Tested | P value<br>(Wilcoxon Rank Sum Test) | Rank | Benjamini-Hochberg<br>significance |
| --- | --- | --- | --- |
| 12 | 0.008113 | 1 | Significant |
| 2 | 0.00931 | 2 | Significant |
| 6 | 0.02766 | 3 | Not significant |
| 4 | 0.02889 | 4 | Not significant |
| 7 | 0.05429 | 5 | Not significant |
| 3 | 0.07053 | 6 | Not significant |
| 13 | 0.07955 | 7 | Not significant |
| 5 | 0.1739 | 8 | Not significant |
| 10 | 0.2184 | 9 | Not significant |
| 1 | 0.2361 | 10 | Not significant |
| 8 | 0.4047 | 11 | Not significant |
| 11 | 0.5407 | 12 | Not significant |
| 9 | 0.6404 | 13 | Not significant |
| 14 | 0.7077 | 14 | Not significant |

**Figure 1:**
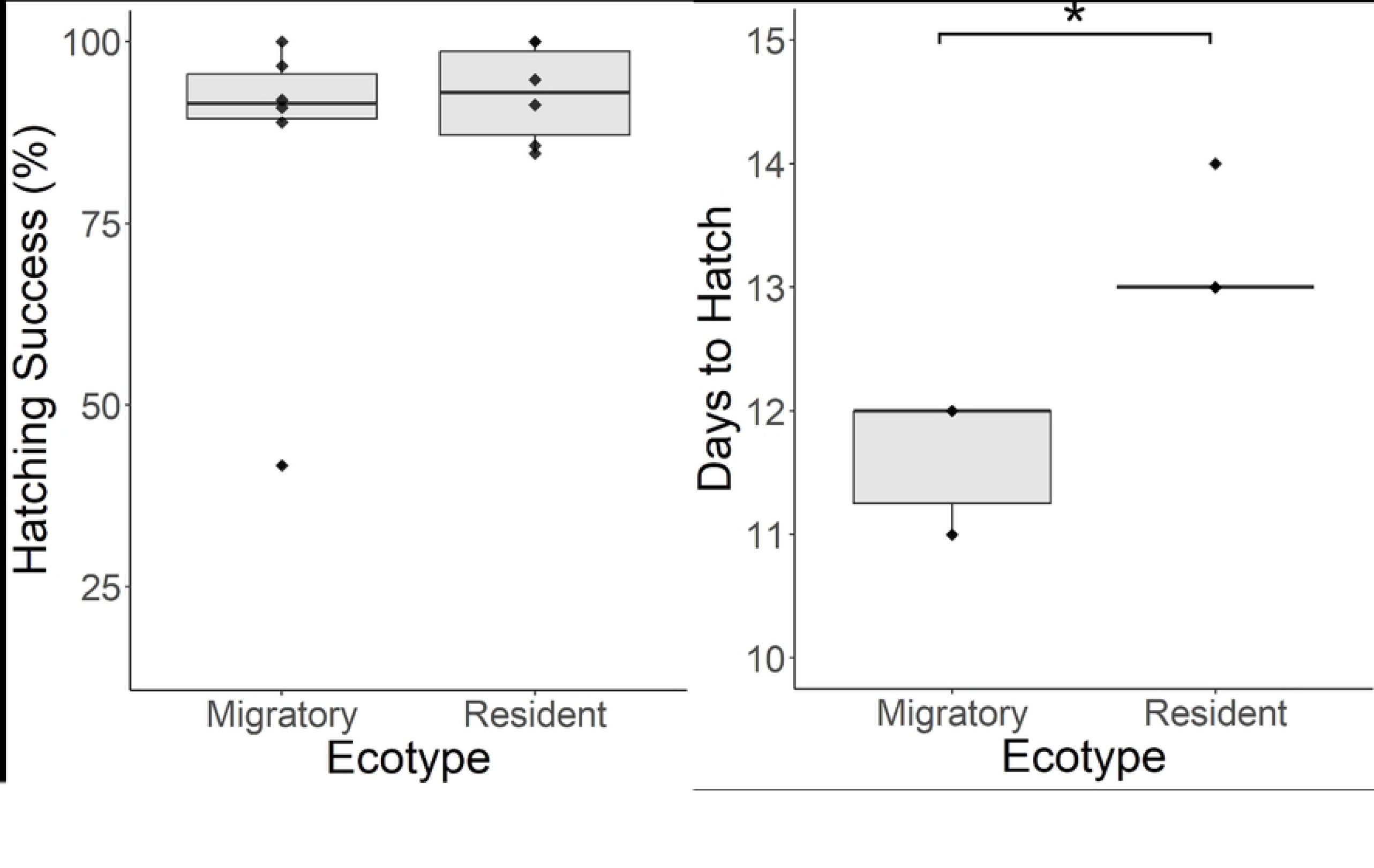
Hatching success (a) and days to hatch (b) for migratory and resident three-spined stickleback incubated at 14°C. Hatching success was calculated per clutch as number of successfully hatched fry divided by initial number of eggs. Days to hatch is calculated from fertilisation (day 0) until the first fry was fully hatched out of the egg. Boxplot shows median and interquartile range and all data values are displayed as points. n = 6 clutches per ecotype. Significant differences, p < 0.001, marked with an asterisk.

Although there were quantitative differences in the timing of development between the ecotypes (see below), qualitatively development was very similar in migratory and resident clutches and therefore we only describe the development of lagoon resident clutches in detail, at each 24 hour interval. Day 0 is the day of fertilisation, day 1 refers to 24-hours post fertilisation, and so on. Figures 2 and 3 show a photo time-series of development which gives a clearer impression than the line drawings that were previously available.

**Figure 2:**
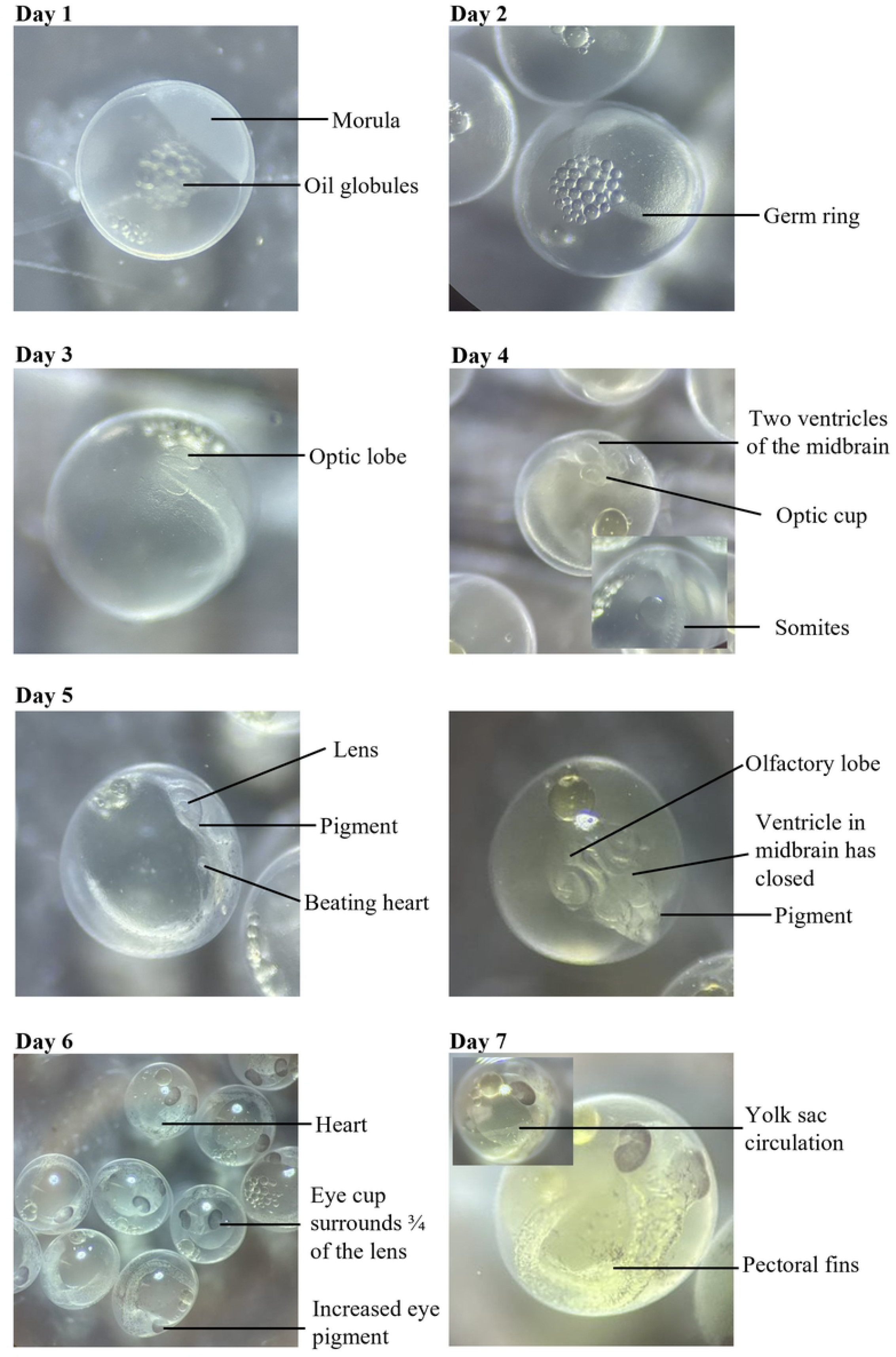
Three-spined stickleback development from day 1 (24-hours post fertilisation) to day 7. Images taken using a dissection microscope with a dark background on the stage, for contrast. Key features are labelled.

**Figure 3:**
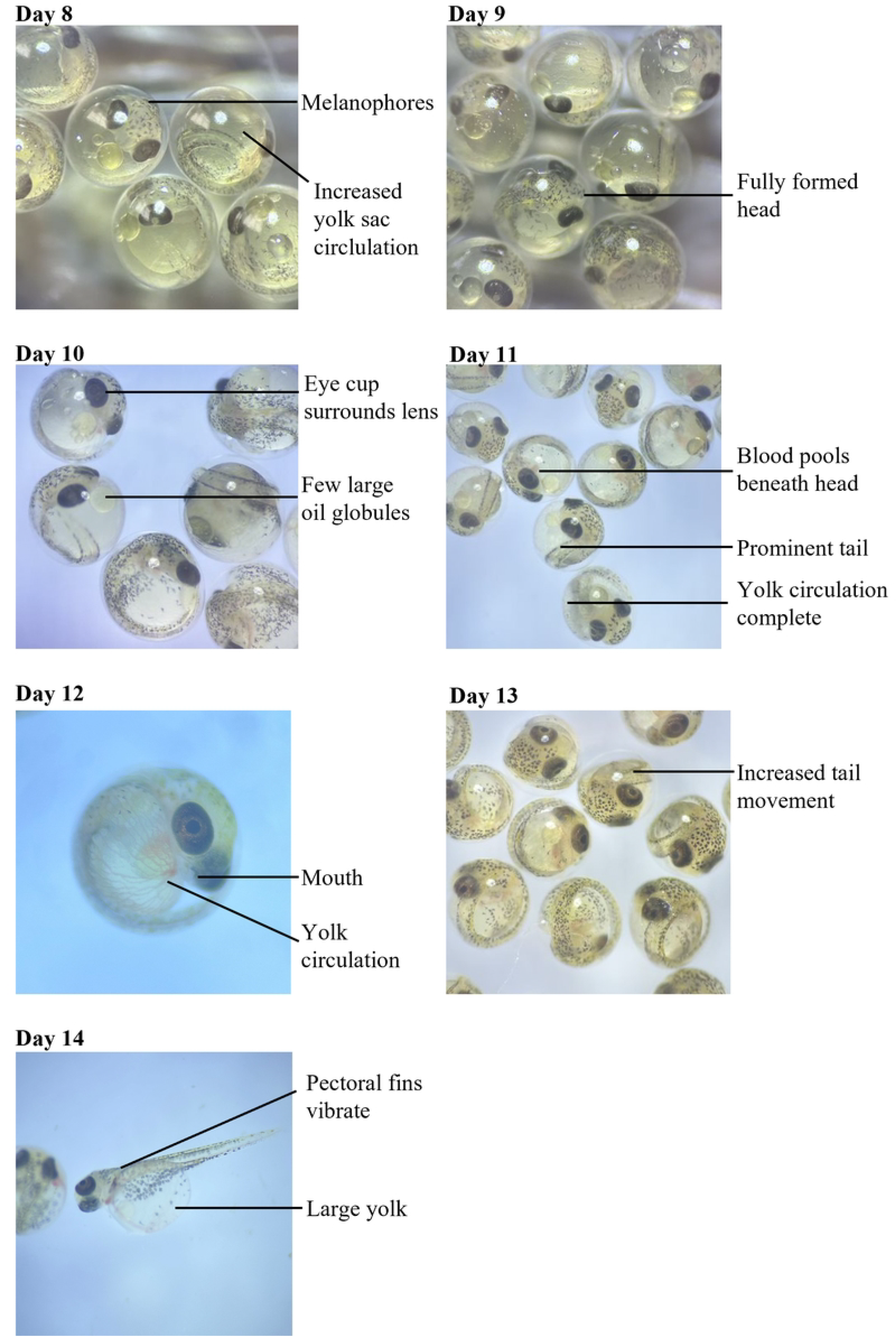
Three-spined stickleback development from day 8 up until hatching. Images taken using a dissection microscope with either a dark or light background on the stage, for contrast. Key features are labelled.

*Day 1:* By 24-hours post fertilisation, the egg has gone through the 2, 4, 8, 16 and 32-cell stage, reaching stage 9 in the Swarup series. The cells divide further, becoming smaller; these form the morula which sits on the yolk. Oil globules are visible in the yolk.

*Day 2:* Gastrulation has begun. A germ ring forms. The more opaque ring of cells seen in Swarup stage 12 were often not visible until the eggs were gently rotated, as the germ ring was usually on the ventral side. A dark background aided visualisation of features early in development.

*Day 3:* Optic lobes have formed either side of the forebrain (Swarup stage 16). The embryo now protrudes from the surface of the yolk sac

*Day 4:* Somites have appeared in the middle of the embryo. Rather than from the side of the embryo, as shown in Swarup 1958 stage 17, somites were best seen by gradually focusing through the yolk sac to the opposite side (Figure 2, day 4). Around six or seven pairs were clear in most cases, although occasionally there were more. The head differentiates so that the optic lobes become optic vesicles where central cavities are visible, and then optic cups as lenses form. The brain develops further so that separation between the mid- and hindbrain is clear.

*Day 5:* The heartbeat is now apparent on the left side of the embryo (Swarup stage 19). This is most clear to see when looking at the embryo from the side, where only one eye is visible (Figure 2, day 5). Some pigment begins to appear, starting on the outer margins of the eye and some scattered areas on the body. Ventricles in the midbrain have closed. Some gentle rotation of the eggs may be necessary to see both the heartbeat and the features of the head, hence two images are provided for day 5 in Figure 2.

*Day 6:* More eye and body pigment is present so that most of the eye cup is dark in colour and this now surrounds almost all of the lens except for the lower portion. The tail occasionally moves, although at this point it was infrequent and difficult to capture. The split in the hindbrain can no longer be seen. The heart is larger now and the three chambers become visible.

*Day 7:* Yolk sac circulation can be seen on the left of the embryo; often gently rotating the embryo is necessary to identify this. Pectoral fins start to develop. There is now more pigment, including on the yolk sac. The head of the embryo becomes shorter and broader (Swarup stage 22).

*Day 8:* More melanophores are visible on the head. The yolk sac circulation increases in area; at this stage this can be difficult to see as the colour is similar to the yolk sac so switching between a dark and light background can help distinguish this. Tail movement becomes more frequent and the eye pigment is darker throughout the eye cup. The split in the forebrain is still visible.

*Day 9:* The ventricle of the forebrain has closed so the head is now fully formed. The yolk sac circulation is almost complete (Swarup stage 23).

*Day 10:* The eye cup now completely surrounds the lens, which becomes dark in colour. The eye cup surrounding the lens happens at an earlier stage than in Swarup’s descriptions as here this occurs prior to the completion of the yolk sac circulation and formation of the mouth, whereas this is noted at stage 24 of Swarup’s descriptions where the embryo is almost ready to hatch (the final stage, approx. 24 hours prior to hatching). There are now only a few larger oil globules and these are situated in front of the head. Blood also collects in front of the head. A change to a lighter background at this point in development was found to increase contrast between the pigmented features and the background, making identification easier.

*Day 11:* The blood collecting in front of the head becomes more apparent as the yolk circulation is now complete and its red colour is clear, especially on a light background. The mouth forms and the tail becomes more prominent. To observe the mouth, gently moving the embryos was necessary to position the head of the embryo in the best light to capture the mouth; it was best observed looking at the front of the embryo (as in Figure 2, day 12).

*Day 12:* The amount of pigment on the body has increased further. Tail movement becomes more frequent and the pectoral fins vibrate. The embryos now display all key features described in stage 24 of Swarup’s taxonomy and are ready for hatching.

*Day 13 or 14:* At day 13 or 14 hatching commences. The head of the embryo pushes against the shell of the egg and breaks free. Tail movement then frees the rest of the embryo. The hatched embryo is transparent and lays on its side as the yolk is large. The head remains curved around the edge of the yolk.

There were differences in development between lagoon resident and migratory stickleback: comparing development each day, migratory clutches were at a significantly higher score than residents at days 2, 4, 6 and 12 (Figure 4, table 1), but after correcting for multiple testing only differences at days 2 and 12 remained significant (table 1). Differences were therefore early in development when migratory clutches had a higher score and were therefore more developed than residents, and late in development (day 12) when the eggs of residents took longer to hatch than migratory clutches after reaching the final stage (24) in Swarup’s scoring (Figure 4). Progression of development in our clutches of migratory and resident stickleback (raised at 14°C) also differed strongly from those of Swarup raised at 18-19°C. Early in development (days 1 to 6) there is only a small difference, with those raised at 18-19°C having, in general, a slightly higher development score (Figure 4). However, by day 7 there is a large difference in score, with Swarup’s having reached stage 24, the final stage before hatching, but both resident and migratory clutches raised at 14°C averaging a score of 22.6 (migratory mean was 22.8 and resident 22.4). Those raised at the higher temperatures hatch (stage 25) at days 6 to 8, while clutches raised at the lower temperature remain at the later stages of development (stages 22 to 24) for a longer time, before hatching between days 11 and 14.

**Figure 4:**
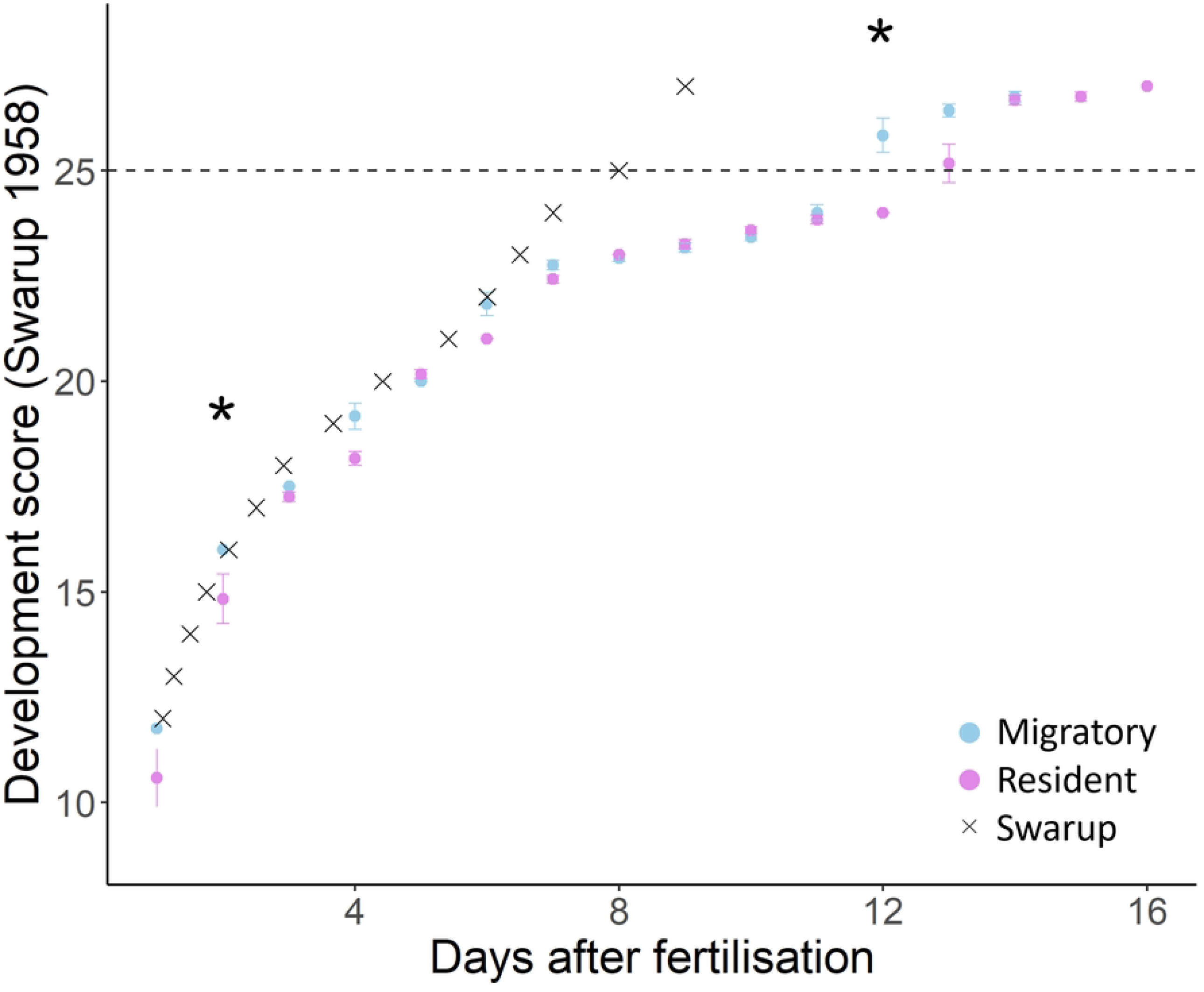
Development of stickleback embryos through time (days after fertilisation) estimated as the Swarup (1958) score. Migratory stickleback clutches are shown as blue points and residents as pink points. Clutches were raised at 14°C. Grey crosses show the development score from Swarup 1958 for comparison, where stickleback were raised at 18-19°C. Dotted line shows when stickleback have hatched. Asterix shows days where there was a significant difference (p < 0.05) between migratory and resident scores (two-sample Wilcoxon rank sum tests) that remained significant after correcting for multiple comparisons using the FDR method: FDR = 0.10.

## Discussion

We describe visual features of development in the embryos of two contrasting ecotypes of three-spined stickleback, every 24 hours post-fertilisation, with key structures highlighted in photographs to increase the ease of identification of important features. This builds upon the work of Swarup (1958), where stages were named after key morphological characteristics and described in detail in stickleback embryos raised at 18 to 19°C. However, as stickleback development is strongly influenced by temperature, this work assessed these key features daily in stickleback developing at a lower, physiologically relevant temperature for where these populations would be found naturally, and to compare the two ecotypes. The hatching success of both ecotypes was high, with no differences between them. We have presented only coloured images of resident stickleback because, aside from the difference in hatching time, qualitatively we did not observe any differences in development between ecotypes. As we viewed embryos every 24 hours it remains possible that finer differences in detail exist but could not be observed here.

Migratory clutches tended to be further developed than residents from days 1 to 7, although this only reached significance at day 2. From day 8, no difference in development stage was observed between ecotypes, so resident clutches had ‘caught up’ with migratory, and both ecotypes had reached stage 24 at the same time. Migratory clutches then hatched on day 11 or 12 while residents remained at the last stage prior to hatching. The eggs of resident stickleback therefore hatched on average 1.5 days later than those of migratory fish. As embryos were checked every 24-hours, the hatch day was recorded as the day the first fully hatched fry was observed. This could therefore be slightly longer than actual time to hatching (as fry may have hatched sometime within the day prior), but this should not cause a systematic bias between the ecotypes. Indeed, the difference between ecotypes was large, with no overlap in hatch day between groups.

The migratory and resident forms are known to vary greatly in morphology and genetics, so a difference in development is not unexpected [6]. As migratory stickleback have larger clutches than resident and there is a trade-off between clutch size and egg size [10], the eggs of migratory fish are smaller than those of residents. The decreased yolk size and therefore nutrients for migratory embryos may explain the reduced time to hatch, as well as additional differences in the egg or membrane itself. Further studies on the egg of both ecotypes would be required to fully understand these findings. Temperature, pH and dissolved oxygen have also previously been implicated in stickleback growth and development (Lefébure et al., 2011; Wanzenböck et al., 2022, Candolin et al., 2022), but as rearing conditions were consistent here, these cannot explain the differences in development between ecotypes.

In addition to the ecotype differences observed, by comparing resident and migratory stickleback raised at 14°C to stickleback raised at 18-19°C by Swarup (1958), the effect of incubation temperature can be explored, although this is limited as Swarup provides only an estimate for the time to reach each stage. All stickleback were raised in freshwater but different populations were used between studies, so additional population effects can also not be ruled out. Time to hatch was greater than 3 days longer at 14°C than Swarup found at the warmer temperature of 18-19°C, but differences in development between temperatures did not become apparent until 7 days post-fertilisation. Temperature therefore has limited influence early on in stickleback development, up until yolk sac circulation, but warmer temperatures then induce hatching earlier. Similar findings have been made in the zebrafish (*Danio rerio*), where the same rate of early development was observed at a wide range of temperatures [13]. In the three-spined stickleback, temperature greatly effects paternal care behaviour, reproductive success and growth rate post-hatching [12,14], but effects on embryo development have rarely been studied.

This study has revisualised the key characteristics in three-spined stickleback development up until hatching, following the landmark work of Swarup (1958). Using North Uist lagoon resident and migratory populations that live in sympatry during the breeding season provides further information on how different ecotypes develop at a temperature they would naturally experience. It provides a set of features that are clear at each day post fertilisation to allow comparison between future treatments, particularly important if we want to discover how stickleback embryos will cope with the increasing water temperature and carbon dioxide levels that may be experienced. These stages also provide a baseline for any abnormalities in development to be compared to. A high yet consistent hatching success between both ecotypes has also been shown and that migratory stickleback hatch at least a day earlier on average than resident stickleback from the same loch. Further investigation into this in natural populations, and studies of migratory and resident stickleback eggs, would be necessary to explain this pattern. However, as this species is often raised in aquaria at the temperature used here, this information adds to our knowledge of how long stickleback incubation lasts before hatching.

## Acknowledgements

We are grateful to Laura Dean, Ann Lowe and Henry Lewis for their help and advice with this project during fieldwork.

